# Cervical gene delivery of the antimicrobial peptide, Human β-defensin (HBD)-3, in a mouse model of ascending infection-related preterm birth

**DOI:** 10.1101/643171

**Authors:** Natalie Suff, Rajvinder Karda, Juan Antinao Diaz, Joanne Ng, Julien Baruteau, Dany Perocheau, Peter W. Taylor, Dagmar Alber, Suzanne M.K. Buckley, Mona Bajaj-Elliott, Simon N. Waddington, Donald Peebles

## Abstract

Approximately 40% of preterm births are preceded by microbial invasion of the intrauterine space: ascent from the vagina is the most common pathway. Within the cervical canal, antimicrobial peptides and proteins (AMPs) help to constitute a barrier which prevents ascending infection. We investigated whether expression of the AMP, human β-defensin-3 (HBD3), in the cervical mucosa prevented bacterial ascent from the vagina into the uterine cavity of pregnant mice. An adeno-associated virus vector containing both the *HBD3* gene and *GFP* transgene (AAV8 HBD3.GFP) or control (AAV8 GFP), was administered intravaginally into E13.5 pregnant mice. Ascending infection was induced at E16.5 using bioluminescent *E.coli* (*E.coli* K1 A192PP-lux2). Bioluminescence imaging showed bacterial ascent into the uterine cavity, cellular events that led to premature delivery and a reduction in pups born alive, compared with uninfected controls. In addition, a significant reduction in uterine bioluminescence in the AAV8 HBD3.GFP-treated mice was observed 24 hours *post-E.coli* infection, compared to AAV8 GFP treated mice, signifying reduced bacterial ascent in AAV8 HBD3.GFP-treated mice. There was also an increase in the number of living pups in AAV HBD3.GFP-treated mice. We propose that HBD3 may be considered a possible candidate for augmenting cervical innate immunity to prevent ascending infection-related preterm birth.

## Introduction

Preterm birth, defined as delivery before 37 completed weeks gestation, affects 11% of pregnancies worldwide^1^. It is the single largest cause of mortality in infants under 5 years old and it is associated with serious morbidity in the surviving infants, particularly for those born before 32 weeks gestation^2^. Prematurity accounts for 29% of global neonatal deaths per year and for 3.1% of total disability adjusted life years in the global burden of disease^3^. Despite extensive research, the rates of preterm birth have remained stable over the years; this is thought to be largely due to a lack of effective preventative treatments.

Preterm birth is a highly complex, multifactorial biological process which culminates in the premature activation of the common parturition pathway^4^. Evidence indicates a role for infection and inflammation in preterm birth, particularly in those occurring before 28 weeks, and it is estimated to be associated with up to 40% of preterm deliveries^5^. In clinical studies, the increased prevalence and diversity of intrauterine bacterial DNA is associated with preterm pre-labour rupture of membranes and spontaneous preterm birth^6^. Bacteria linked with preterm birth include those genera associated with relatively low pathogenicity such as *Ureaplasma, Fusibacterium, Mycoplasma* and *Streptococcus*^6^. Once in the pregnant uterus, bacteria interact with the mucosal lining and the local immune system initiating an inflammatory cascade leading to cervical ripening and myometrial contractility, ultimately resulting in preterm parturition^7^. Animal models confirm this link; inoculation of the intrauterine cavity with live bacteria or bacterial pattern recognition pattern (PRR) moieties, such as lipopolysaccharide (LPS), leads to preterm birth ^8,9^. In humans, this preterm inflammatory pathway has proved resistant to most therapies including antibiotics, cervical cerclage and tocolytics; meta-data analysis shows that progesterone appears to delay preterm birth, although its role in infection and inflammation remains unclear^10^.

Ascending vaginal infection is considered to be the most common route by which bacteria gain access into the uterine cavity in cases of spontaneous preterm birth^11^. This hypothesis is supported by the association recorded between the bacterial species identified in the amniotic fluid, fetal membranes and placenta and those normally found in the lower genital tract ^6,12^. Additionally, recent data suggests that dominance of a particular bacterial species within the vaginal microbiota are associated with an increased risk of preterm birth^13^.

The ability of the human cervix to prevent bacteria ascending from the vagina to the uterine cavity depends on several factors, including a mucus plug that provides a negatively-charged platform for interaction with cationic antimicrobial peptides (AMPs) including the Human β-defensins (HBDs)^14,15^. It is not known why certain women develop ascending infection; women who have had previous cervical excisional treatment are associated with an increased risk of preterm birth^16^, but it is likely that an endogenous compromise in cervical mucosal immunity plays a key role in these cases^17^. In support of this, recent evidence has shown an association between PTB and low mid-trimester cervico-vaginal levels of human β-defensin 2^18^.

HBDs, produced by the cervical epithelium, critical for maintaining mucosal host-microbial homeostasis^19^. In addition to their direct microbicidal role against pathogens, they mediate inflammation by influencing cytokine production, immune cell chemotaxis and epithelial cell proliferation^20^. Human β-defensin 3 (HBD3) has potent and broad-spectrum antimicrobial activity against bacteria, fungi and viruses^21,22^. Possessing multiple positively charged arginine and lysine residues it has the highest net positive charge of all AMPs which probably contributes to its broad-spectrum action. Its antimicrobial activity is not affected by differing physiological salt concentrations, making it an ideal choice for a clinical treatment^21^. Furthermore, unlike other β-defensins, HBD3 binds to bacterial products, such as LPS, resulting in reduced pro-inflammatory cytokine responses^23^.

The transfer and expression of therapeutic genes in the context of cervical mucosa has been explored as a potential treatment for infectious diseases. For example, anti-HIV antibodies have been delivered using an adeno-associated viral vector (AAV) to human cervical and vaginal cells *in vitro* and also to the lower genital tract of rhesus macaques to successfully prevent mucosal acquisition of HIV infection *in vivo*^24,25^. Gene transfer to the murine cervix, however, has not been described before, although the use of adenoviral vectors to deliver specific pro-inflammatory cytokines to the mouse vagina has been explored as a possible treatment for vaginal candidiasis^26^.

The objectives of our present study were to evaluate the function of the HBD3 gene, delivered to the cervix using AAV8, in a mouse model of ascending infection-related preterm birth. We have previously established an ascending infection-model of preterm birth using bioluminescent *E.coli* K1 A192PP-*lux2* (*E.coli* K1)^27,28^, a strain known to cause neonatal sepsis and meningitis in rats. In this model, *E.coli K1* administration induced delivery significantly earlier than in pregnant mice receiving intravaginal PBS as well as leading to a significant reduction in the proportion of pups born alive compared with intravaginal PBS controls^28^.

Here, we tested the hypothesis that cervical gene expression of HBD3 reduces microbial ascension into the pregnant uterine cavity, reducing the frequency of preterm birth.

## Results

### A murine model of ascending bacterial infection and preterm birth

The murine model of infection was essentially as described previously by our laboratory^28^. The ability of a pathogenic strain of *E.coli* K1 A192PP-*lux2* (*E.coli* K1) to ascend into the embryonic day 16.5 pregnant uterine cavity was investigated^28^. Bioluminescence imaging of the dam revealed ascent of bacteria to the top of the uterine cavity by 24 hours (Fig.1A), with diffuse spread of bacteria in the fetal membranes, the placenta and the amniotic fluid by 18 hours (Fig.1B). By 24 hours bacteria were evident in the fetus (Fig.1C); immunoperoxidase staining for bacteria revealed microbial presence within the respiratory and gastro-intestinal (GI) tract (Fig.1D). By postnatal day 1 (approximately 72 hours post-infection), bacteria were predominantly seen in the GI tract (Fig.1E).

**Figure 1.**
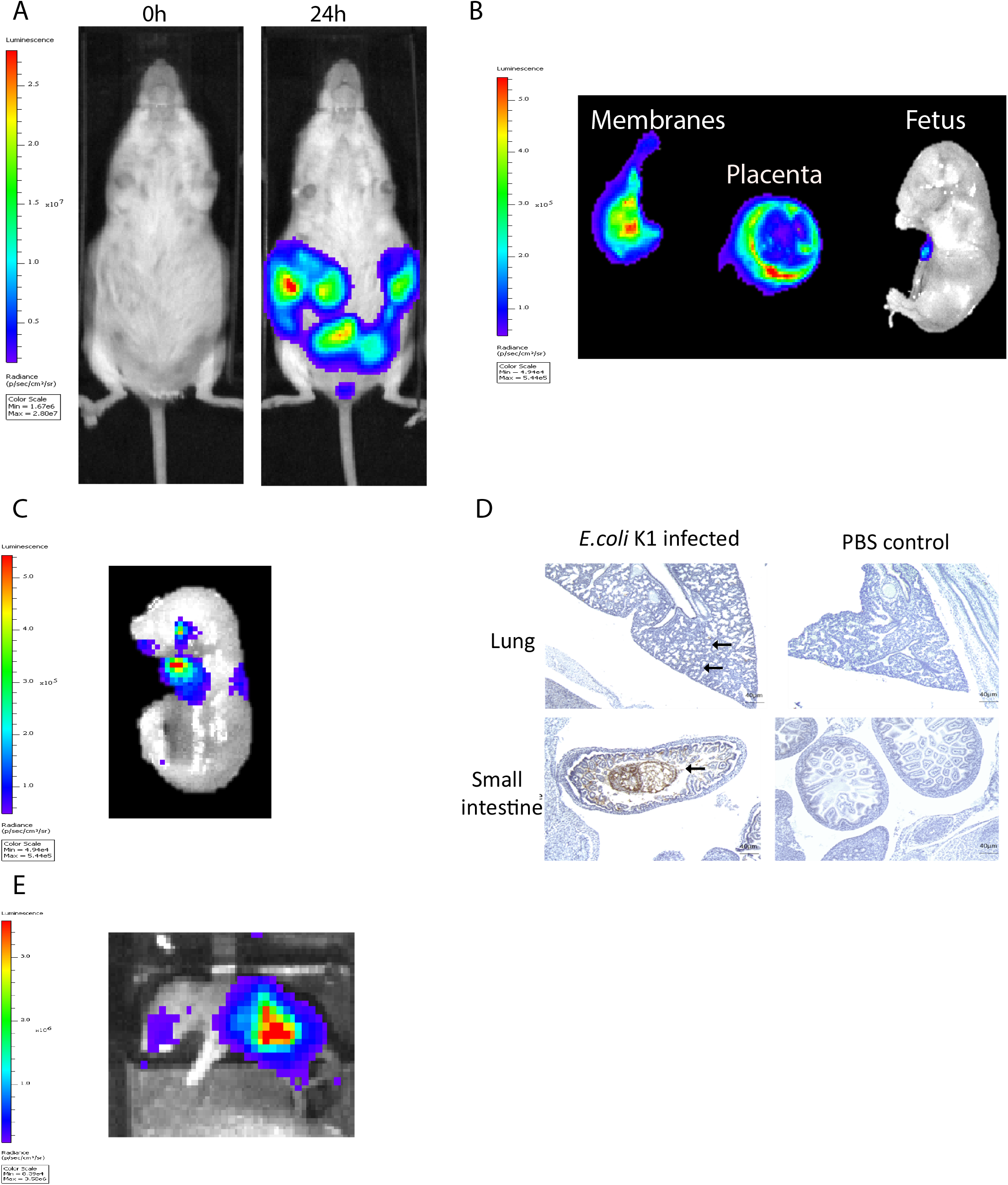
Intravaginal bioluminescent *E.coli* K1 A192PP-lux2 (*E.coli* K1) can ascend into the pregnant uterine cavity and induce premature delivery (30). (A) Pregnant mice received intravaginal *E.coli* K1 on embryonic day 16.5, bacteria ascend into the uterine cavity over 24 hours. (B) At 18 hours after *E.coli* K1 administration bacteria is seen specifically within the pregnant uterine cavity and is detected in the fetal membranes, the placenta and the amniotic fluid by 18 hours. (C) By 24 hours, bacteria is detected in the fetus. (D) Immunohistochemical detection of *E.coli* in the fetus shows E.coli specifically within the lung alveoli and small intestine at 24 hours, whilst no *E.coli* is seen in the uninfected fetus (sections counterstained with haematoxylin), n=3, Scale bar 40μm. (E) By Postnatal day 1 (72 hours after infection), *E.coli* is clearly seen in the Gastrointestinal tract.

### Selection of viral vector for optimal gene transfer to the cervical mucosa

To determine if viral vectors are capable of delivering genes to the cervical mucosa, separate cohorts of non-pregnant adult mice received intravaginal delivery of either adeno-associated virus serotype 2/8 (AAV8), recombinant adenovirus serotype 5 (rAD5) and VSVg pseudotyped HIV lentivirus, each containing the firefly luciferase transgene. Luciferase expression was seen in the lower genital tract 48 hours to 120 hours after vector administration (Fig. 2A). Although there was high luciferase activity from VsVG pseudo-typed HIV lentivirus, AAV was chosen for the ensuing studies due to its relatively low immunogenicity, episomal-nature, thermostability and success in numerous clinical trials^29–31^. Following AAV8 administration, it was confined to the upper vagina and cervix at 72 hours with no spread to the uterus or liver (Fig. 2B and Fig.2C). This vector was used for the remaining experiments due to its relatively low immunogenicity and success in a wide range of pre-clinical and clinical trials^29,32^.

**Figure 2.**
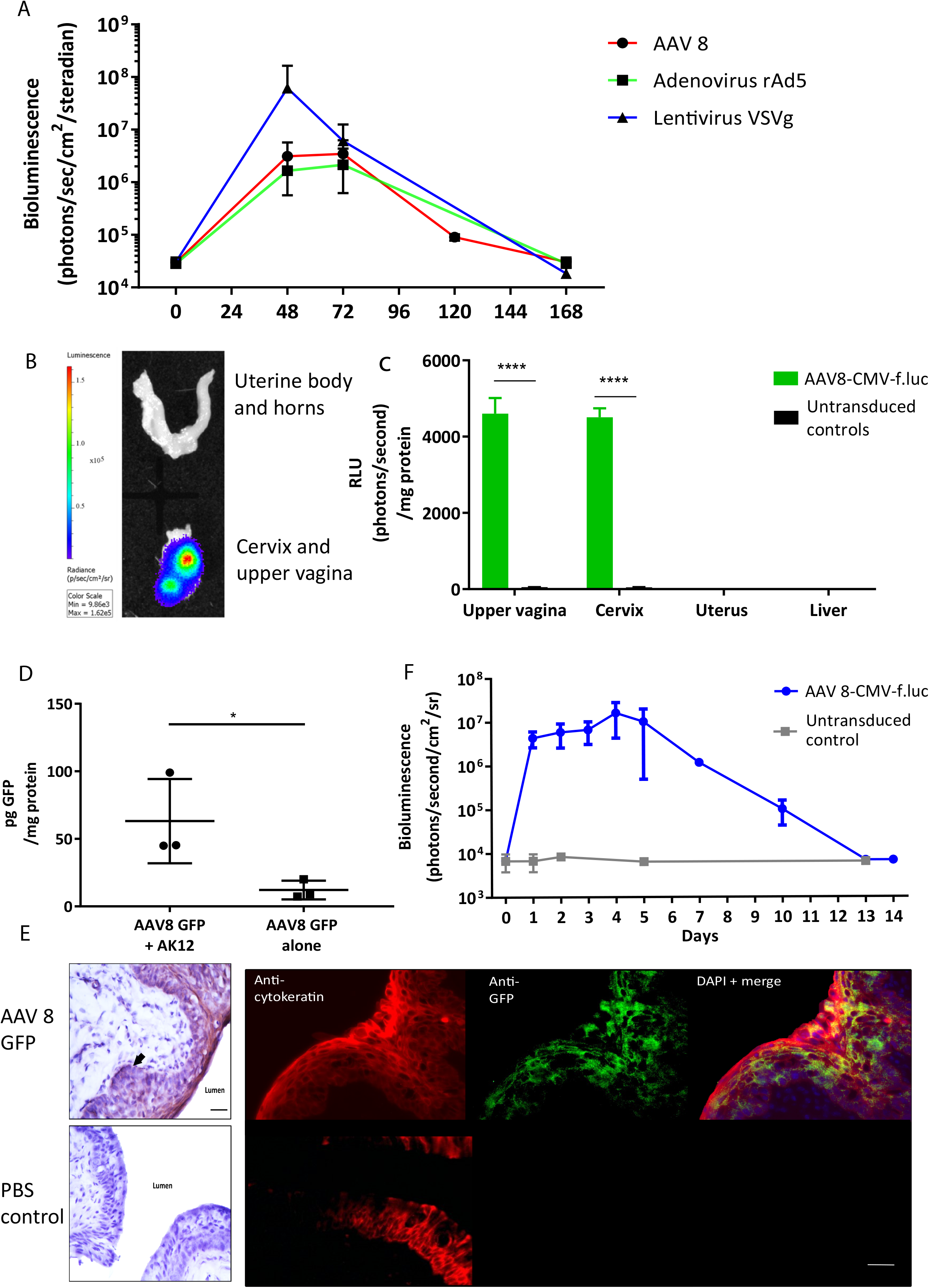
Gene delivery to the cervix is possible using viral vectors. (A) Adenovirus-associated virus-8 (AAV-8), Recombinant adenovirus 5 (rAd5), VSVg lentiviral vectors can deliver luciferase to the cervix resulting in transient luciferase expression. (B, C) 72 hours following AAV8-CMV-f.luc administration, luciferase expression is limited to the cervix and upper vagina, n=3. ****P<0.0001, data were analyzed by a 1-way ANOVA with post-hoc Bonferroni test. Gene transfer to the cervix using AAV 8 viral vector is improved when delivered with the pluronic gel, AK12, and results in transient protein expression in the epithelial cell layers. (D) Cervical GFP expression is increased when AAV8 GFP is delivered in combination with AK12, compared with AAV8 GFP delivery alone, n=3. *P<0.05, data were analyzed by a unpaired T-test. (E) GFP expression is detected in the epithelial layers of the cervix, confirmed by protein co-localisation with cytokeratin expression. Scale bar 20μm. (F) Following delivery of AAV8-CMV-f.luc in combination with AK12 gel, luciferase expression in the cervix lasted for up to 14 days, with peak expression occurring between day 3 and day 5, n=3.

### Specific location of vaginal and cervical protein expression

Thermosensitive pluronic gels have been developed for numerous functions, including vaginal drug delivery (26–28). We assessed AK12, that gels at 30 degrees Celsius, as a method to prolong the contact time of the vector with the epithelium to improve transduction. Delivering AAV8 GFP in combination with this gel intravaginally resulted in significantly higher GFP expression than delivering vector alone (*P=0.02*, Fig.2D). This expression was co-localised with cytokeratin, a marker of the cervical epithelial cell layers (Fig.2E). Following AAV-8 luciferase intravaginal administration bioluminescence imaging revealed luciferase expression to behighest between 3 and 5 days after transduction and lastingfor up to 14 days (Fig. 2F).

### HBD3 expression and function

Following intravaginal administration of AAV8 HBD3.GFP, HBD3 and GFP expression was co-localised in the cervical and upper vaginal epithelium after 72 hours (Fig.3A). HBD3 was detected in the vaginal lavage 96 hours after vector administration, indicating that the HBD3 peptide was appropriately synthesised and secreted into the mouse cervix and vagina (Fig.3B).

**Figure 3.**
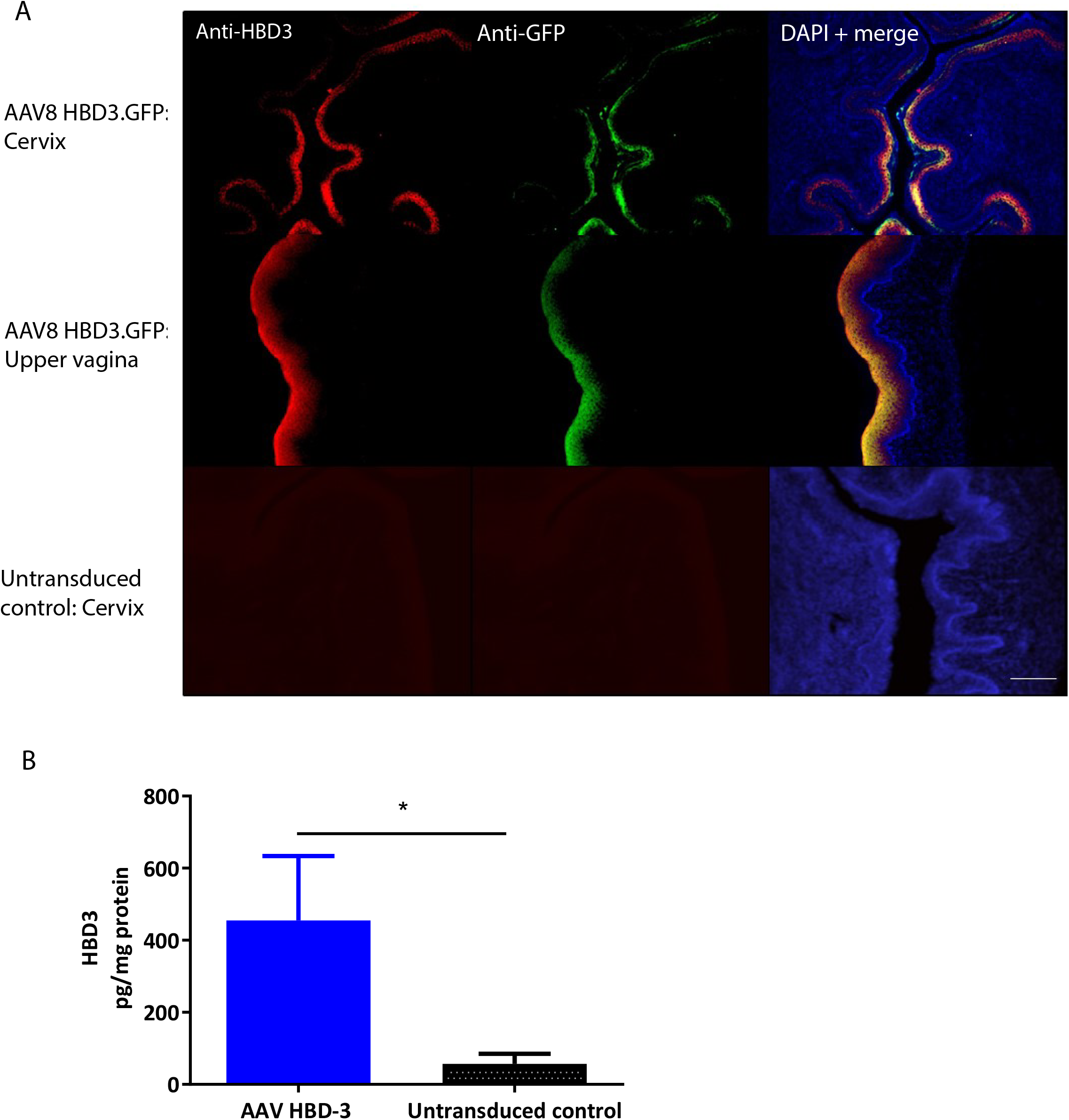
Cervical delivery of the AAV8 HBD3.GFP vector results in HBD3 peptide expression in the epithelial layers of the cervix and upper vagina and is secreted into the vaginal fluid. (A) Upper vaginal and cervical tissue were harvested 72 hours after vector or PBS administration. HBD3 and GFP are detected in the cervix and vagina following AAV HBD3.GFP administration, n=3. Scale bar 20μm. (B) Vaginal lavage was collected from a different cohort of mice 96 hours after administration of the vector or PBS, n=3. *P<0.05, data were analyzed by an unpaired T-test.

We performed *E.coli* killing assays with these lavages. A trend for bactericidal activity was observed in AAV HBD3.GFP treated *versus* PBS controls, but this did not reach statistical significance (Fig.4A). Immunohistochemical detection of neutrophils was performed on cervical tissue 72 hours after AAV.HBD3 GFP, AAV.GFP and PBS intravaginal administration (Fig.4B). This showed a significant increase in neutrophil recruitment to the cervical epithelium layer in the AAV8 HBD3.GFP group 72 hours after vector administration, compared with the AAV8 GFP and PBS only controls (*P=0.009* and *P=0.017*, respectively, Fig.4C).

**Figure 4.**
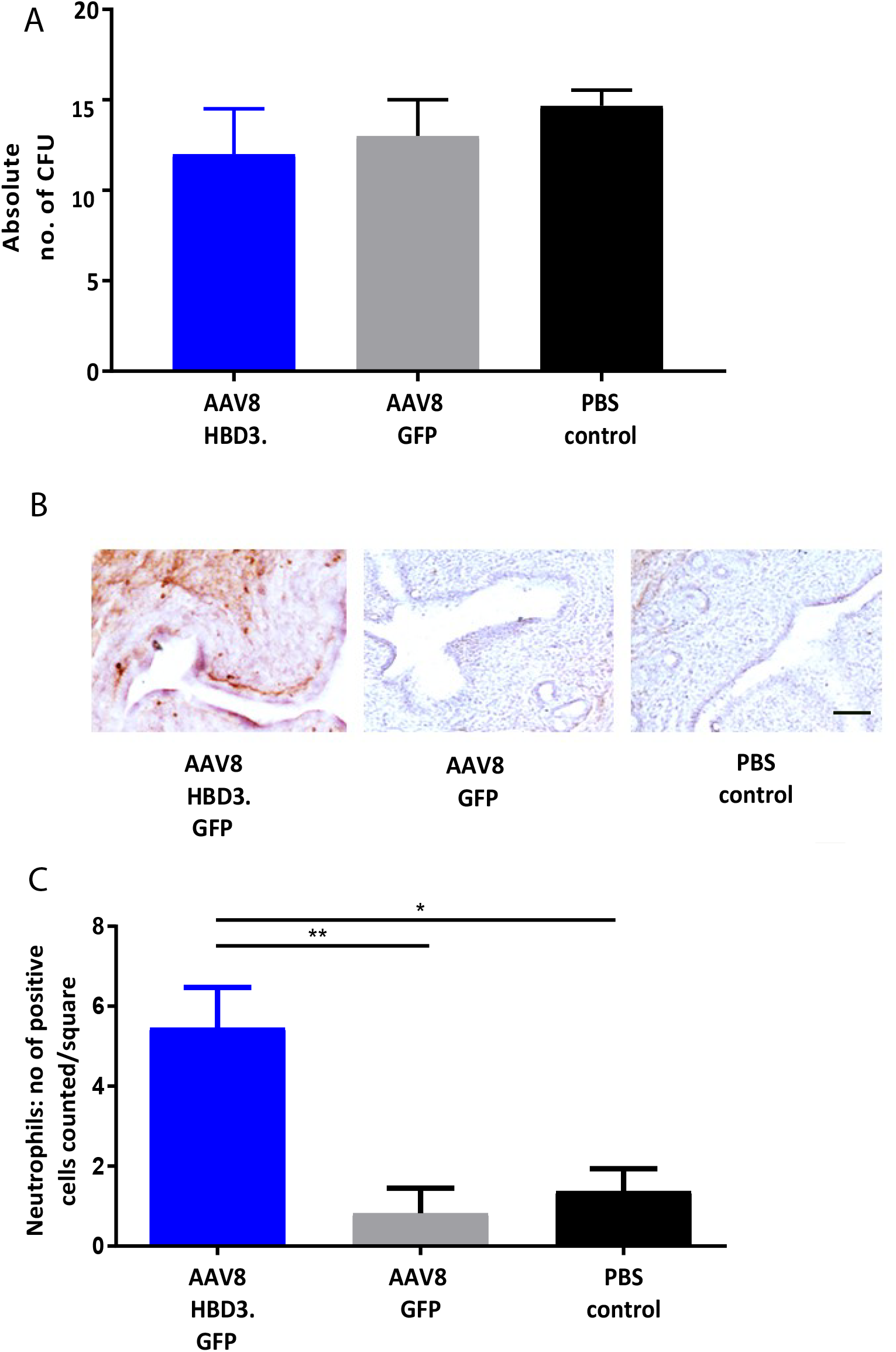
AAV8 HBD3.GFP increases neutrophil recruitment to the upper cervical epithelial layers. (A) *E. coli* killing assays were performed on vaginal lavages from AAV8 HBD3.GFP, AAV8 GFP and PBS treated mice. There was no difference in bacterial kill, n=3. Data were analysed with a 1-way ANOVA and post-hoc Bonferroni tests. (B) Representative images of cervical neutrophil localization using immunohistochemical staining of Ly-6g with haematoxylin counterstain. Brown coloration depicts DAB-positive cells, Scale bar 20μm. (C) Neutrophil numbers in cervical epithelial cell layers were increased following AAV8 HBD-3 transduction, compared with AAV8 GFP and PBS, n=3. *P<0.05, **P<0.005, data were analysed with a 1-way ANOVA and post-hoc Bonferroni tests (

Vaginal lavage samples were taken before- and at multiple time points after-vector or PBS administration to investigate the effect of HBD3 on the vaginal microbiome. Delivery of AAV8 HBD3.GFP had no effect on the alpha diversity index (Supplementary Fig.1A) or the distribution of bacterial classes compared to the AAV8 GFP control group was similar before and after vector administration (Supplementary Fig.1B).

### HBD3 gene delivery to prevent ascending infection

Next we investigated the effects of AAV8 HBD3.GFP delivery on ascending infection and preterm birth. AAV8 HBD3.GFP or AAV8 GFP control was administered to the vaginas of pregnant dams on embryonic day 13.5, followed by administration of *E.coli* K1 intravaginally on embryonic day 16.5. Representative images of an AAV8 HBD3.GFP treated and AAV8 GFP control mouse are shown in Fig.5A. AAV8 HBD3.GFP administration resulted in significantly less uterine bacterial bioluminescence, a marker of bacterial ascent, at embryonic day 17.5 (*P=0.0015*). A similar trend at embryonic day 18.5 did not reach statistical significance (*P=0.09*) (Fig.5B).

**Figure 5.**
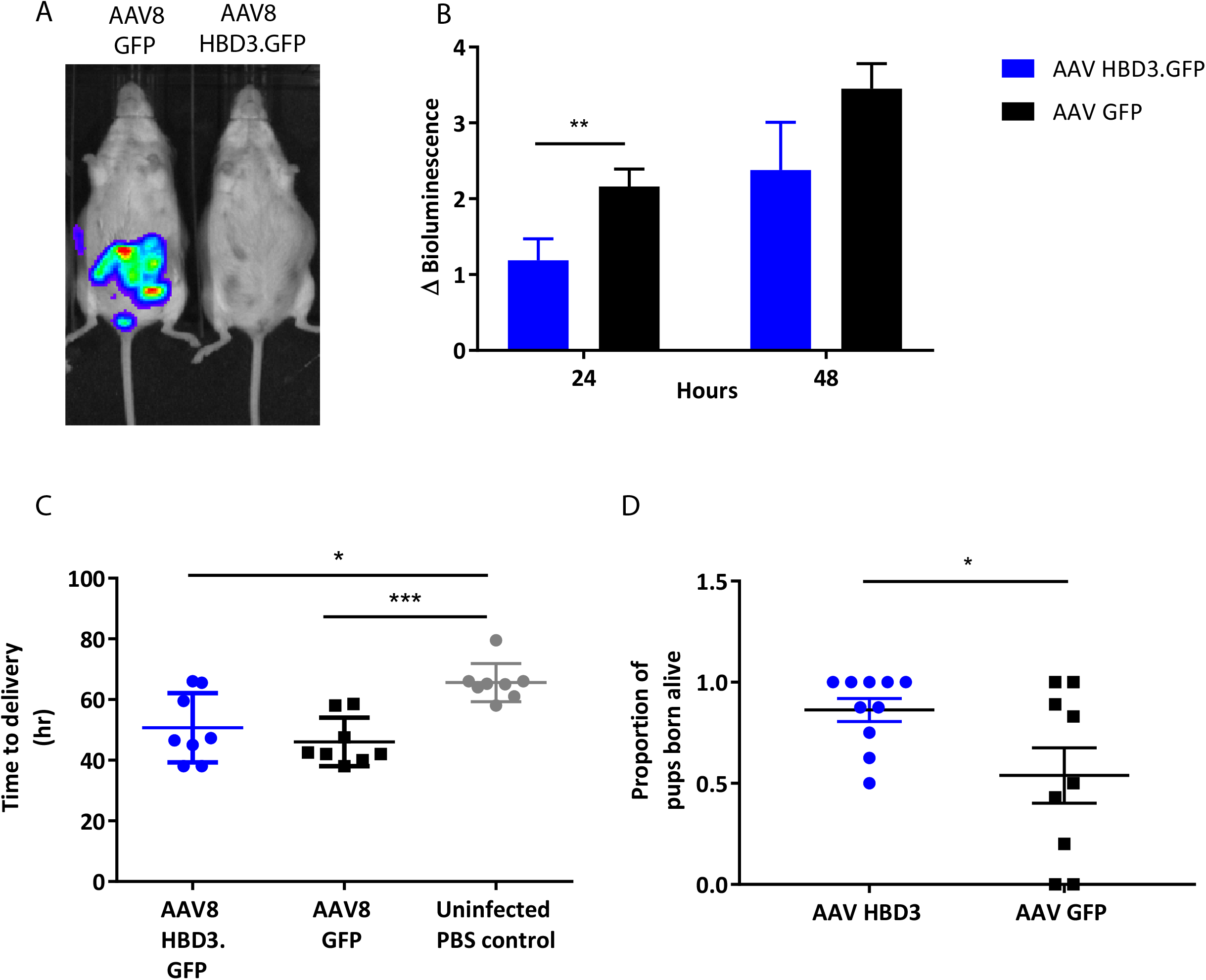
Delivery of AAV8 HBD3.GFP reduces bacterial ascent into the pregnant uterine cavity and increases the proportion of pups born alive. (A) An example of an AAV8 HBD3.GFP treated mouse and an AAV8 GFP treated mouse 24 hours after bacterial administration. (B) Marked difference in uterine bacterial bioluminescence in mice treated with AAV8 HBD3.GFP, compared with AAV8 GFP, n=10. **P<0.01, data were log-transformed and analysed with a repeated measures 2-way ANOVA with post-hoc Bonferroni tests. (C) The AAV8 HBD3.GFP group delivered significantly earlier than the control mice not infected with bacteria and the mean time of delivery was not significantly different fromthat of the AAV8 GFP group n=10. *P<0.05, ***P<0.001, data were log transformed and analysed with a 1-way ANOVA and post hoc Bonferroni tests. (D) There was an increase in the proportion of pups born alive in the AAV8 HBD3.GFP treated group, compared with the AAV8 GFP group, n=10. *P<0.05, data were arc-sin transformed and analysed with an unpaired test.

The reduced ascent of bacteria into the uterine cavity in the AAV8 HBD3.GFP group led us to hypothesise that this group would also have reduced preterm birth rates. Delivery within 48 hours (embryonic day 18.5) of intravaginal administration of *E.coli* K1 was considered preterm birth, whereas mice delivering after this point was considered term. AAV8 HBD3.GFP resulted in a small, nonsignificant reduction in preterm labour to 60% versus 78% in AAV8 GFP controls (*P=0.37*, Table 1).

**Table 1.**
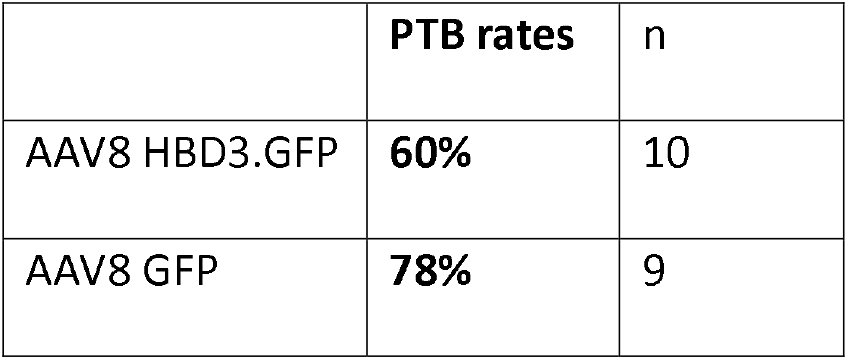
Ascending infection-induced preterm birth rates: Preterm birth rates after intravaginal administration of *E.coli* K1 on embryonic day 16.5. Data analysed with Fisher exact test.

The AAV8 HBD3.GFP group delivered significantly earlier than the control mice not infected with bacteria (mean 50.7h ± 11.4 vs. 65.5h ± 6.3, *P=0.01*, Fig.5C) and the mean time of delivery was not significantly different from that of the AAV8 GFP group (mean 50.7h ± 11.4 vs. 46.1 ± 8.0, *P=0.95*, Fig 5C).

AAV HBD3.GFP treatment significantly increased the proportions of live pups, versus AAV8 GFP controls (0.86 vs 0.54 respectively (Fig.5D), Unpaired t-test; *P*=0.028) but did not increase survival rates at seven days (AAV8 HBD3.GFP 60.3% vs AAV8 GFP 55.9%, *P*=0.2) (Fig.6A). However, there was a small non-significant increase in survival in those pups born at term in the AAV8 HBD3.GFP group (85.2% vs. 62.5%, respectively, *P*=0.16, Fig.6B).

**Figure 6.**
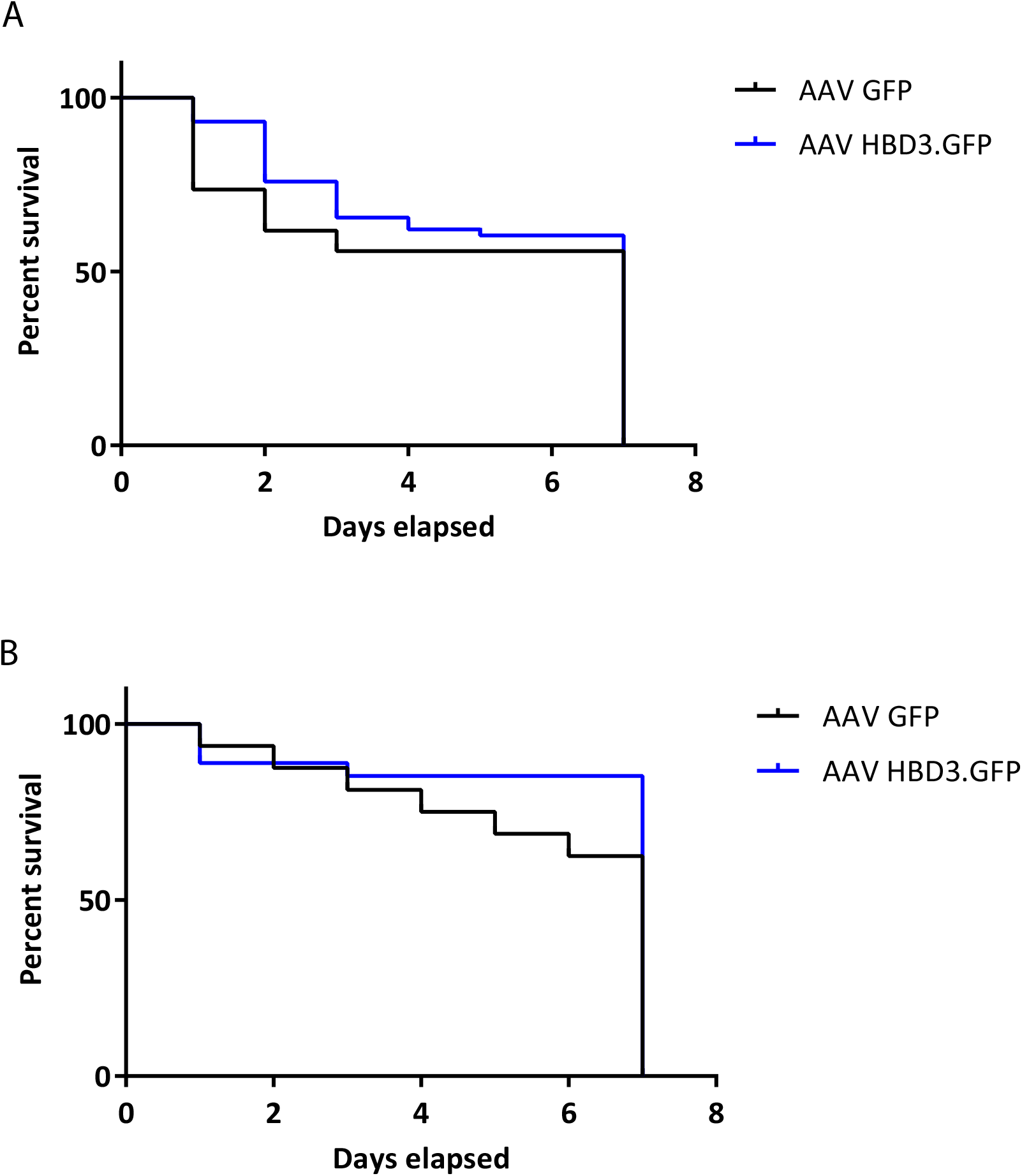
There is no difference in pup survival over the first week of life between AAV HBD3.GFP and AAV GFP pups, although there is a trend for increased survival in the term AAV HBD3.GFP pups compared with AAV GFP controls pups. (A) Overall pup survival over the first week of life. (B) Term pup survival over the first week of life. Term pups were defined as those that were delivered > 48 hours after bacterial administration. n=53 in the AAV HBD3.GFP group, n=27 in AAV GFP group (Term pups; AAV HBD3.GFP n= 23, AAV GFP n=10). Data analysed by Log-rank Mantel-Cox test.

## Discussion

Despite recent advances in preterm birth research, current therapies do not appear to have an impact on preterm birth rates^36^. Approximately 40% of preterm births are thought to be associated with infection, specifically ascending infection from the vagina and despite this knowledge, prophylactic antibiotics have not be shown to alter preterm birth rates^37^. Here, we show that a localised cervical treatment, which delivers an endogenous human gene with antimicrobial properties, can reduce ascending infection in pregnant mice and lead to an increase in neonatal survival.

We show that gene delivery to the murine cervix is possible using common viral vectors and is augmented by the use of thermosensitive pluronic gel. In contrast to the human cervix, the entirety of the murine cervix comprises stratified squamous epithelium^38^. Although little is known about the mouse cervix, we do know that one layer of the 28-layer human vaginal and ectocervical stratified squamous epithelium sheds every four hours^39^ and so pluronic gels may improve transduction efficiencies by increasing contact time of vector with epithelium. Pluronic gels have previously been used in combination with adenovirus vector to deliver VEGF to the uterine circulation as a treatment for fetal growth restriction^40^. The benefits of these gels are that they are biodegradable, low in toxicity and they transfer from aqueous phase to gel phase on increasing temperature. We chose the AK12 gel for our experiments as its gelling temperature of 30°C means that it forms a gel quickly upon intravaginal application preventing vector spillage and it also seemed appropriate for a plug that would be exposed to the cooler outside environment at the vaginal introitus. Furthermore, there is evidence that pluronic gels help to facilitate cervico-mucus plug penetration without comprising the function of the barrier^41^.

AAV8 HBD3.GFP did not significantly increase the gestational length in *E.coli* K1 infected dams. However, there was an increase in the proportion of pups born alive from dams in this group with a tendency towards increased 7 day survival rates for neonates born at term following *E.coli* K1 infection. There was no increase in preterm pup survival in the first week of life, however, suggesting that pups delivered early are more susceptible to *E.coli* K1 infection.

The mechanisms by which cervical HBD3 reduces bacterial ascent into the uterine cavity is uncertain. Although HBD3 is known for its potent antimicrobial action, we were unable to measure a statistical difference in the microbicidal activity of the vaginal lavage from AAV8 HBD3.GFP *versus* AAV8 GFP treated mice in the experimental time frame. Interestingly, within the same time frame, the presence of HBD3 resulted in an increase in neutrophil influx into the cervical epithelium *in vivo*. Therefore, it is likely that HBD3-mediated bacterial clearance involves activation of multiple antimicrobial mechanism(s). A study looking at the benefits of exogenously delivered human cathelicidin LL-37 in a murine model of *P.aeruginosa* lung infection found that cathelicidin LL-37 enhanced bacterial clearance *in vivo^42^*. Interestingly, the authors found no evidence of a direct microbicidal effect and it was concluded that the likely mechanism of bacterial clearance is peptide-mediated enhanced early neutrophil influx. Our observations also support a crucial role for HBD3-driven neutrophil influx, taken together, it is tempting to suggest that neutrophil mediation may present a significant armoury of AMP action.

HBD-3 concentrations detected in the murine cervicovaginal lavage following gene transfer was lower than compared with physiological levels in human cervicovaginal lavage fluid (0.5ng/mg protein versus 55.5ng/mg protein)^43^. This may be primarily be due to species variation and/or due to the experimental setup of our current study e.g cervicovaginal samples were taken 72 hours following gene transfer whilst bioluminescence (indicative of HBD3 gene expression) continued to increase up to 96 hours post-gene delivery (Fig 2A). Future experiments, exploring dose-response and time course of expression will be valuable for providing further insight into HBD3 production. Cervicovaginal levels of endogenous mouse defensin proteins were not assessed, however, our unpublished data suggests a trend towards reduced *Defb14* (the gene for mBD14, the mouse orthologue of HBD-3) mRNA levels in the AAV8 HBD-3 treated cervices raising the hypothesis that HBD-3 is involved in a negative feedback loop suppressing *Defb14* expression.

Recent data has shown that human cervical mucus plugs contain at least 28 AMPs, although interestingly, the AMPs were at insufficient concentrations to have antimicrobial activity against Group *B streptococcus*, a bacteria commonly implicated in preterm birth^44^. However, these AMP appear to have a role in amplifying the immune response including enhanced leucocyte activity and complement-mediated killing. In support of this, emerging evidence suggests that HBDs have a much more complex role in the immune system than being solely endogenous antimicrobials, they have previously been shown to recruit immune cells, such as neutrophils, macrophages and dendritic cells^45,46^. In contrast to our findings here, HBD-3 has previously been reported to have direct chemotactic activity on human monocytes and not neutrophils^46^. Our unpublished data showed minimal cervical monocyte influx. The chemotactic activity on murine neutrophils seen in this study, may be due to species variation and its mechanism, direct or indirect, needs to be elucidated in future studies. Neutrophils are critical effector cells and form part of the first line of defence against microorganisms. HBD2 has been shown to attract neutrophils via the G-protein phospholipase C-dependent pathway^45^. The CCR6 receptor is involved in activation of this pathway; HBD3 and mouse β-defensin 14 are chemotactic for mouse CCR6-expressing HEK 293 cells *in vitro^47,48^*. Furthermore, the cysteine residues of the three disulphide bridge structure of HBD3 appear to be necessary for this chemotactic role^49^. In addition to its chemotactic role, HBD3 has been shown to prolong the lifespan of neutrophils at sites of infection by preventing apoptosis^50^. The molecular mechanism(s) involved in HBD3-mediated neutrophil influx in our model system require further investigation.

During human pregnancy, a rise in oestrogen levels leads to a stable vaginal microbiome dominated by *Lactobacillus* species^51^. *Lactobacillus* are thought to inhibit pathogen growth by secreting antimicrobial bacteriocins as well as producing lactic acid which helps to maintain a low, acidic pH^52^. The composition of the vaginal microbiota appears to be closely associated with preterm birth outcome^13,53^. Mice treated with either AAV8 HBD3.GFP or AAV8 GFP did not show significant changes in their vaginal microbiota nor did it modify *Lactobacillus* species abundance in the vagina (data not shown). In support of these findings, HBD3 does not exhibit antibacterial activity against lactic acid bacteria, including *L.rhamnosus*, suggesting that these bacteria may exhibit factors that protect them from host immune defence^54^. Recent data indicates an association between an *L.iners*- dominant vaginal microbiome and an increased risk of preterm birth, whilst L.crispatus-dominance appears to be protective^13^. *L.crispatus* promotes epithelial cell defence against *Candida albicans in vitro* by increasing HBD levels, in particular, HBD3^55^. This finding may play a role in linking the mechanism of Lcrispotus-dominant vaginal microbiome with its protective effect on preterm birth.

HBD3 was previously considered to be pro-inflammatory as its expression increases following TLR activation or IL-1β, TNF-α and IFN-⍰ release^21,46^, however, emerging data recognises it more as a multifunctional effector of the immune system^56^. It is possible that HBD3 may induce a strong chemoattractant and inflammatory response at high concentrations during infection and injury to respond to the insult whilst at lower concentrations it has anti-inflammatory and healing properties^57^. In this study, we found no evidence of increased inflammatory cytokine expression in the cervical tissue following gene transfer of AAV.HBD3 GFP compared with AAV GFP (Supplementary Fig.3). Further studies would need to be conducted to assess for other surrogate markers of inflammation or cytotoxicity. In the GI tract, HBD3 plays an important role in contributing to mucosal immune tolerance^58^. As the vaginal microbial composition appears to be so closely related to preterm birth risk, HBD3 expression in the cervix may improve immune tolerance to certain commensal bacteria.

The use of antimicrobial gene therapy has been explored previously, mainly as a way of treating infections with antibiotic resistant bacteria and treating certain wound infections where the skin epithelium has been severely damaged, such as burns ^59^. Gene delivery of HBD3 to keratinocytes using adenoviral vectors has been performed in studies investigating novel treatments for wound infections ^60,61^. They show that keratinocytes expressing HBD3 *in vitro* have increased antimicrobial activity and interestingly found that the HBDs worked synergistically with the AMP, cathelicidin LL-37, in its antimicrobial activity against *E.coli* and *S. aureus*^61^. In an *in vivo* porcine model of infected diabetic wounds, adenoviral vectors delivering HBD3 resulted in reduced bacterial load 4 days after vector administration and an improvement in re-epithelialization of the wound^60^. The wound healing role of HBD3 may also be of interest in women at high risk of preterm birth who have had previous cervical treatment for high risk precancerous cervical lesions. The expression of HBD3, in combination with VEGF, in bone marrow derived stem cells resulted in improved wound healing in a model of combined radiation-wound injury in rats^62^. In particular, HBD3 is shown to promote skin re-epithelialization, granulation tissue formation and collagen I deposition. In preterm birth, premature remodelling and shortening of the cervix involves degradation of cervical stromal collagen, particularly collagen I and III, by matrix metalloproteinases ^63_65^. Therefore, HBD3 gene therapy could help to inhibit this process by encouraging collagen I deposition.

Preterm birth remains a major global health problem, being responsible for greater than a million neonatal deaths per year^66^. Despite the employment of current preventative strategies for preterm birth, there has been no decline in preterm birth rates highlighting the importance of further research into novel therapies^67^. The use of HBD3 in augmenting cervical mucosal immunity may have a role in reducing the intra-uterine inflammation associated with ascending vaginal infection-related preterm birth and warrants further studies to explore its potential for clinical translation.

## Materials and Methods

### Viral vector production

The following viral vectors were used in this study; AAV2/8, VSV-g pseudotyped lentivirus, recombinant adenovirus serotype 5. The vectors carried the firefly luciferase or enhanced GFP transgene under the control of the cytomegalovirus (CMV) promoter. Lentiviral vectors were gifted from Dr Stephen Howe (University College London, UK). The adenoviral vectors were gifted from Dr Alan Parker (Cardiff University, UK). The AAV viral vectors were developed and purchased from Vector Biolabs (Malvern, USA).

An AAV8 bicistronic vector encapsidating a single-stranded DNA sequence containing the HBD3 gene (Vega Sanger HBD3 transcript VEGA68:CM000670.2) under the transcription activity of the CMV promoter, followed by the eGFP gene under the transcription activity of a further CMV promoter, and BGH polyA downstream from the HBD3 and eGFP gene was generated (Vector Biolabs, Malvern, USA) (Supplementary Figure 2).

### Animals and Treatments

All animal studies were conducted under UK Home Office license 70/8030 and approved by the UCL ethical review committee. C57BL/6N-Tyr^c-Brd^ mice were obtained from the Charles River Laboratory (Oxford, UK) and adult mice (6-12 weeks) were time mated. The following morning (when a vaginal plug was noted) was designated as embryonic day 0.5.

#### Animal injections

Adult (6-8 weeks old) female C57BL/6N-Tyr^c-Brd^ mice were anaesthetised with isoflurane in 100% oxygen. 10μl of virus (diluted in PBS if necessary) to a concentration of 1×10^12^ genomic copies/mL was administered intravaginally using a 200μl sterile pipette tip. This was applied in combination with 20μl of AK12 thermosensitive pluronic gel (PolySciTech, Indiana, U.S.A)

For pregnancy experiments, vector was administered as described above on embryonic day 13.5.

#### Ascending vaginal infection model

The ascending vaginal infection model was developed using *E.coli* K1 A192PP modified to contain the *lux* operon from *Photorhabdus luminescens* (*E.coli* K1 A192PP*-lux2*). 20μl of mid-logarithmic-phase *E.coli* (1×10^2^ *E.coli* K1 A299 resuspended in 10mM phosphate buffer), or 20μl of phosphate-buffered saline (PBS) in control animals, was delivered into the vagina of pregnant mice anaesthetised with isoflurane using a 200-ml pipette tip on embryonic day 16.5.

Following bacterial administration, mice were placed in individual cages and continuously monitored with individual closed-circuit television cameras and a digital video recorder ^68^. Time to delivery was recorded and defined as the number of hours from the time of bacterial administration to delivery of the first pup. The number of live and dead pups were recorded. Living pups were weighed daily and were culled if there was a 10% loss in body weight (in accordance with PPL 70/8030).

#### Whole-body bioluminescence imaging

Adult mice were anaesthetised with isoflurane in 100% oxygen. Neonatal mice (up to postnatal day 6) remained conscious during imaging ^69^. Mice were imaged using a cooled charged-coupled device camera, (IVIS machine, Perkin Elmer, Coventry, UK) for between 1 second and 5 minutes. The regions of interest (ROI) were measured using Living Image Software (Perkin Elmer) and expressed as photons per second per centimetre squared per steradian (photons/second/cm^2^/sr).

### Tissue collection

Non-pregnant mice were sacrificed 72 hours after viral vector administration. Pregnant mice were sacrificed 18 hours and 24 hours after intravaginal infection. Mice were anaesthetised using isoflurane, the right atrium incised, and PBS injected into the left ventricle for exsanguination. Uterine tissue was stored in 4% PFA. Embryos were stored in 10% neutral-buffered formalin. A separate cohort of mice were sacrificed by cervical dislocation and vagina, cervix, uterus, placenta and fetal membranes were collected and stored at −20°c for protein analysis.

### Storage of fixed cervical and vaginal tissues

Cervical and vaginal tissues were stored in 4% PFA for 48 hours, transferred to 30% sucrose at 4°C then 40μm transverse sections obtained using a microtome. Embryos were stored in 10% neutral-buffered formalin for 48 hours, followed by storage in 70% ethanol before paraffin embedding and sectioning at 5μm.

### *Ex vivo* luminometry

Tissue samples were lysed with 500μl of 1× Lysis buffer (Promega) followed by homogenisation. The homogenates were centrifuged for 10 minutes at 18000g and the supernatants collected. Each sample was loaded on to a white 96 well plate. 1.5mM of luciferase (Promega) was added at a 1:1 volume ratio to the sample. A FluoStar Omega microplate reader (BMG labtech) was used to read the luminescence and the results were analysed using MARS data analysis data software (BMG labtech).

### HBD3 enzyme-linked immunosorbent assay (ELISA)

HBD3 concentrations in vaginal lavage was measured by ELISA (PeproTech, London, UK) per manufacturer’s instructions. Results were read using the FluoStar Omega microplate reader and analysed in MARS data analysis data software.

### GFP immunohistochemistry and co-localisation immunofluorescence

Representative sections of the organ were selected, mounted onto double-coated chrome gelatin Superfrost slides (VWR, Leicestershire, UK) and left to dry. The slides were placed in 4% PFA for 10 minutes followed by washing in TBS (1× Tris-buffered saline). They were treated with 30% H_2_O_2_ in TBS for 30 minutes at room temperatures and then blocked with 15% normal goat serum (Vector Laboratories, Peterborough, U.K.) in 0.1% TBS-T (0.1% of Triton X-100 in 1× TBS) for 30 minutes at room temperature. This was followed by incubation in primary anti-GFP antibody (Abcam, Cambridge, U.K.) in 10% normal goat serum in 0.1% TBS-T overnight at 4°C. The slides were washed and secondary antibody (Abcam, Cambridge, U.K.) in 10% serum in 0.1% TBS-T was added for 2 hours at room temperature. Following this the slides were incubated for 2 hours in ABC Vectastain (Vector Labs, Peterborough, UK). Slides were transferred into DAB solution (0.05% 3.3’-diaminobenzidine (DAB) in TBS with 30% H_2_O_2_ and left for 2 to 3 minutes. The slides were air dried, dehydrated in 100% ethanol and placed in Histoclear (National diagnostics, USA), followed by cover slipping with DPX mounting solution.

A similar protocol was followed for co-localisation immunofluorescence staining; slides were incubated in two primary antibodies (in 10% normal goat serum in 0.1% TBS-T); anti-GFP antibody (Abcam, Cambridge UK) and anti-pan cytokeratin antibody (Abcam, Cambridge, UK) or anti-HBD3 antibody (Abcam, Cambridge, UK). This was followed by incubation with two corresponding Alexafluor secondary antibodies (Abcam, Cambridge, U.K.) in 10% normal goat serum in 0.1% TBS-T. Sections were then incubated with DAPI for 2 to 3 minutes and then washed in TBS. The sections were dried away from direct sunlight and then coverslips mounted using Fluoromount-G. Sections were stored at 4°C.

### Neutrophil immunohistochemistry and influx quantification

Neutrophil immunohistochemistry was performed using the same protocol to GFP immunohistochemistry above; with primary rat anti-Ly6g antibody and Goat anti-rat secondary antibody (Abcam, Cambridge, U.K.).

Cervical sections were visualised using a x5 objective lens and the epithelium and sub-epithelial stromal areas were identified. 5 random fields of view were selected using a x40 objective lens. The numbers of neutrophils were counted per area. 5 sections were counted per mouse and the number of neutrophils counted per area averaged per mouse

### *E.coli* immunofluorescence

Formalin-fixed embryos were paraffin embedded and sectioned. Paraffin embedded slides were placed in Histoclear for 10 minutes and then rehydrated in ethanol. Antigen retrieval was then performed by boiling the slides in citrate buffer for 20 minutes followed by washing in 1×TBS. Slides were blocked with 15% normal goat serum in 0.1% TBS-T for 30 minutes at room temperature. This was followed by incubation in primary rabbit anti-*E.coli* antibody (Abcam, Cambridge, U.K.) in 10% normal goat serum in 0.1% TBS-T overnight at 4°C. The slides were washed and secondary Goat anti-rabbit IgG H&L Alexa Fluor^®^ 488 (Abcam, Cambridge, U.K.) antibody in 10% serum/ 0.1% TBS-T was added for 2 hours at room temperature. Following this the slides were incubated for 2 hours in ABC Vectastain (Vector Labs, Peterborough, UK). Slides were transferred into DAB solution (0.05% DAB/30% H2O2/TBS) and left for 2 to 3 minutes. The slides were air dried, dehydrated in 100% ethanol and placed in Histoclear (National diagnostics, USA), followed by cover slipping with DPX mounting solution.

### *E.coli* killing assays

*E.coli* K1 was grown to mid-logarithmic phase and diluted to 1×10^5^ colony forming units(CFU)/ml. The bacteria were centrifuged at 14000g for 3 minutes. The pellet was washed once in 10mM phosphate buffer followed by further centrifugation and re-suspension in 10mM phosphate buffer. In a 96 well plate, 90μl of the resuspended bacteria was mixed with 90μl vaginal lavage or 10mM phosphate buffer. The plate was incubated for 30 minutes at 37°C. 20μl of each sample was then mixed with PBS (to inhibit further AMP activity). Serial dilutions were then plated and placed at 37°C overnight. CFUs were counted the following morning.

### Cervical inflammatory cytokine analyses by quantitative PCR

In a separate cohort of mice; cervices were collected and stored in *RNAlater* at −80°c for quantitative PCR (qPCR) analysis. Total RNA was extracted using the RNeasy mini kit (Qiagen, UK), as per the manufacturer’s guidelines. Total RNA was reverse transcribed with the High Capacity cDNA Reverse Transcription kit (Applied Biosystems, USA). Primer sets were obtained from Life Technologies (Supplementary Table 1) and qPCR was performed in the presence of SYBR green. Target gene expression was normalized for RNA loading by using *GAPDH*, using the 2^−ΔΔCt^ method of analysis. All qPCR analyses were performed on an Applied Biosystems QuantStudio 3 instrument (Applied Biosystems, USA).

### Bacterial DNA extraction

DNA extraction of bacterial DNA from frozen vaginal lavage samples was done using Qiagen spin protocol as per manufacturer’s instructions with an additional bead beating step (Qiagen DNA mini kit, Denmark).

### 16S DNA sequencing

The DNA from the above step was quantified using a Qubit DNA high sensitivity assay kit and Qubit 2.0 machine (Thermo Fisher Scientific, UK). The DNA concentration in each well was normalised to the lowest concentration sample. The DNA was then pooled including negative DNA extraction controls. This library was diluted to 0.4nM after quantification using the Qubit 2.0, standard curve qPCR and an Agilent high sensitivity DNA kit with the Agilent 2200 Tapestation instrument (Agilent genomics, Santa Clara, US). Library preparation was carried out using dual-indexed forward and reverse primers, with barcodes. Library preparation PCR was performed. The resulting amplicon was cleaned and pooled using AMPure XP beads (Beckman Coulter) as per manufacturer’s instructions. Each plate was pooled into an equimolar final library after quantification using a Qubit 2.0 (Life technologies). Library was loaded onto a MiSeq (Illumina) as per manufacturer’s protocol for 500 cycle V2 kits with the addition of custom sequencing primers for read 1, read 2 and index 1. Data was analysed using QIIME software (v1.8.0).

### Statistics

Data are expressed as means ±SEM. Time-to-delivery data were log-transformed before analysis, and the proportion of live born pups was arc-sin transformed before analysis. Data were analyzed by unpaired t-tests, one-way ANOVAs and two-way ANOVAs (with post-hoc Bonferroni tests). All statistical analyses were performed with GraphPad Prism software version 7.0. *P*<0.05 was considered statistically significant.

## Acknowledgments

NS received funding from the Wellbeing of Women Research training fellowship grant RT414 and the Priory foundation. SNW received funding from UK Medical Research Council grants; G1000709 and MR/N026101/1, MR/R015325/1, MR/P026494/1 and MR/N019075/1, and from SPARKS 17UCLOl. RK received funding from grants; MR/P026494/1, SPARKS grant 17UCLOl. JN received funding from grants; MR/K02342X/1, GOSHCCVI284, Rosetrees M576. JB received funding from a Research Training Fellowship from Action Medical Research grant GN2137. DMP receives support from the UCLH NIHR Biomedical Research Centre. *E.coli K1 A192PP-lux2* was derived by virtue of a project grant from Action Medical Research GN2075 awarded to PWT. We thank Dr Grace Logan for the analysis of the vaginal microbiome 16S sequencing data.

## Author contributions

N.S., R.K., J.N., J.B., D.P. and J.A. performed experiments. N.S., S.M.K.B., D.A., M.B., P.W.T., S.N.W. and D.P. designed experiments, analysed, and interpreted data. N.S. wrote the manuscript. S.N.W., D.P., and M.B-E edited the manuscript.

---

**Supplementary Figure 1.**
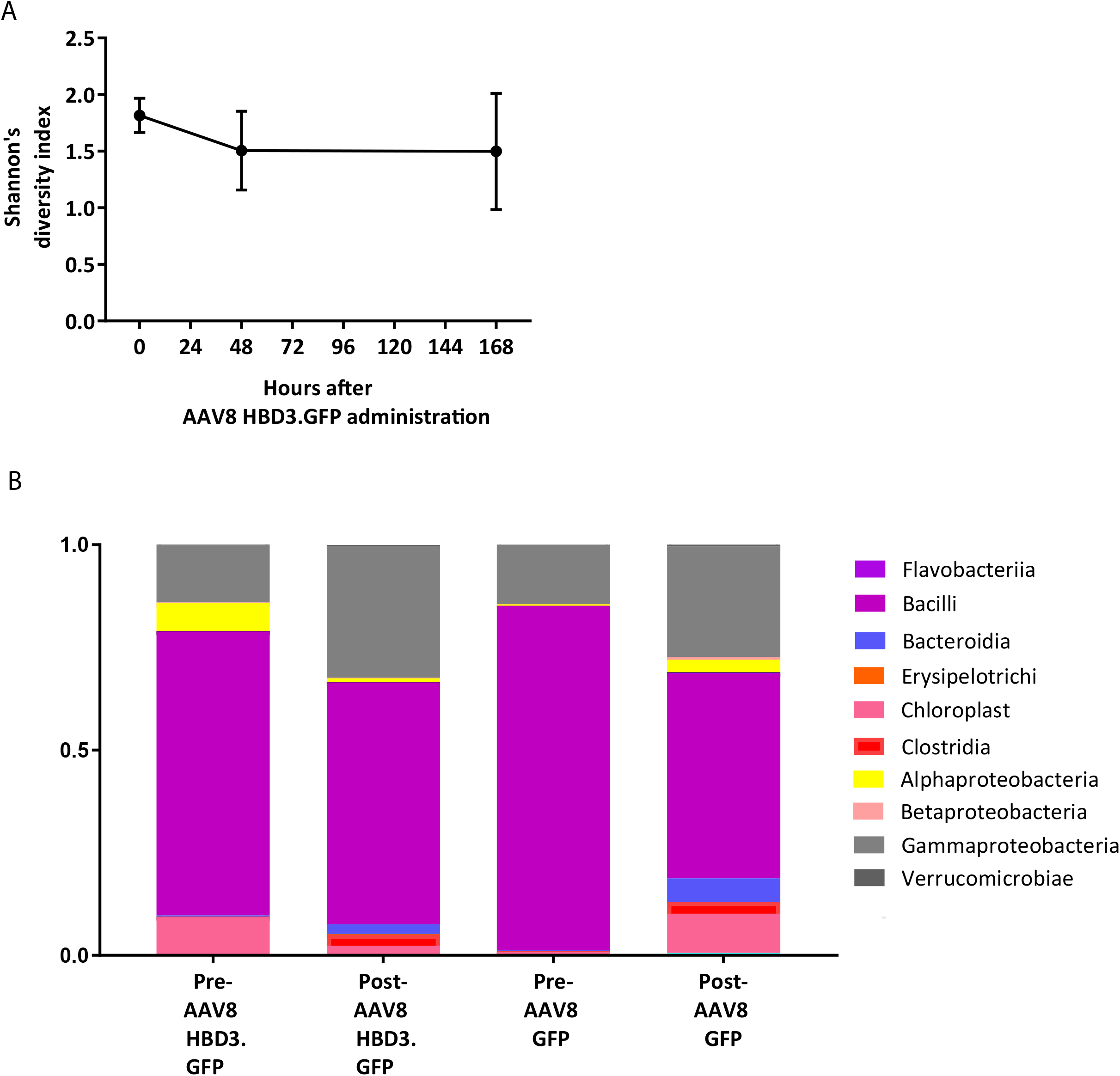
AAV HBD3.GFP does not have an effect on the vaginal microbiome. (A) Alpha diversity was determined before, at 48 hours and at168 hours after AAV8 HBD3.GFP administration, n=5. There was no difference in alpha-diversity values after AAV8 HBD3.GFP, compared with samples taken before AAV8 HBD3 administration. Data were analysed by a 1-way ANOVA with post hoc Bonferroni tests to before AAV8 HBD3.GFP samples. (B) The proportion of bacterial classes present in the vaginal microbiome was determined before and 7 days after AAV8 HBD3.GFP or AAV8 GFP administration, n=5.

**Supplementary Figure 2.**
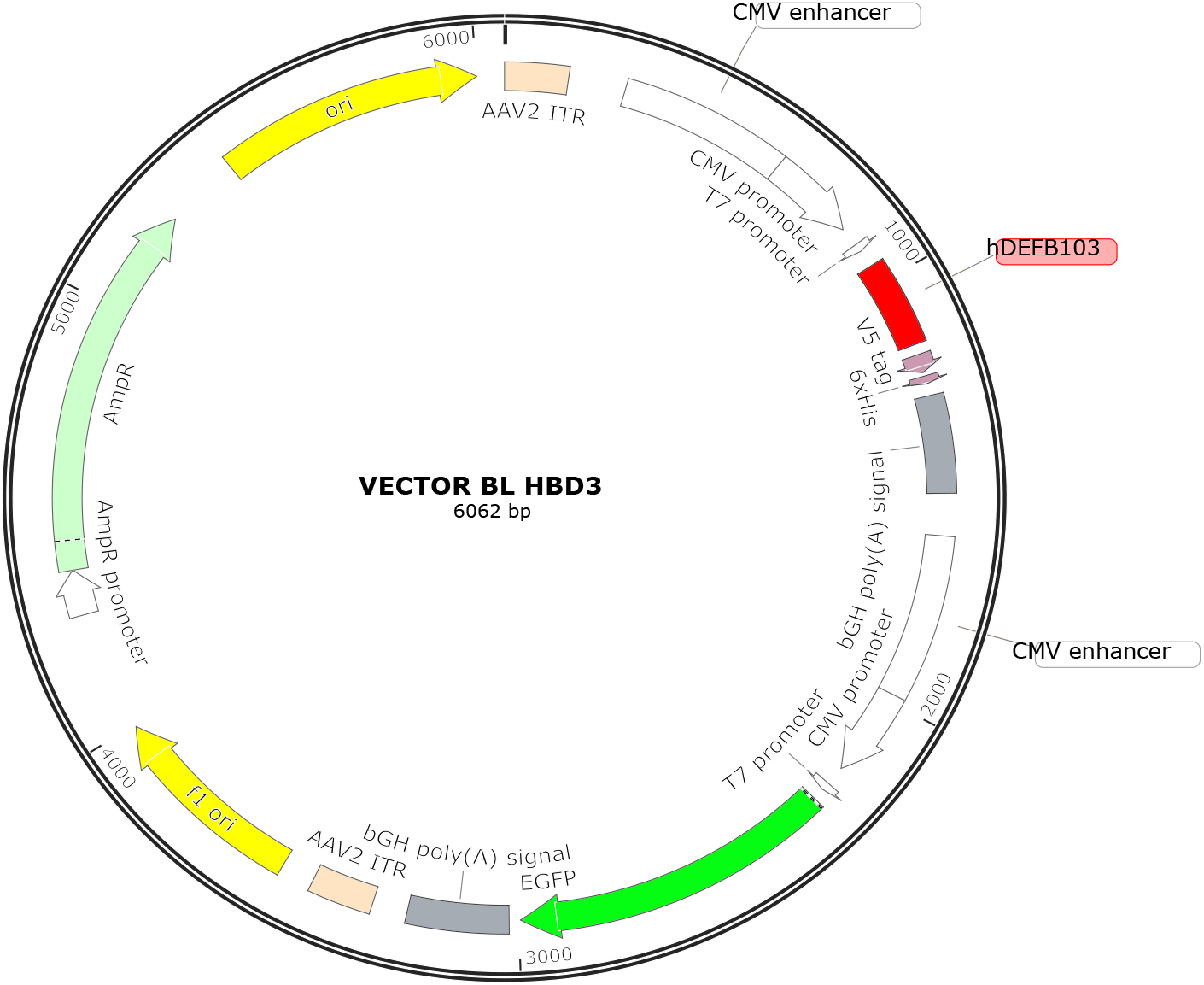
AAV HBD3.GFP construct. An AAV8 bicistronic vector encapsidating a single-stranded DNA sequence containing the HBD3 gene (Vega Sanger HBD3 transcript VEGA68:CM000670.2) under the transcription activity of the CMV promoter, followed by the eGFP gene under the transcription activity of a further CMV promoter, and BGH polyA downstream (Vector Biolabs, Malvern, USA) (Supplementary Figure 2).”

**Supplementary Figure 3.**
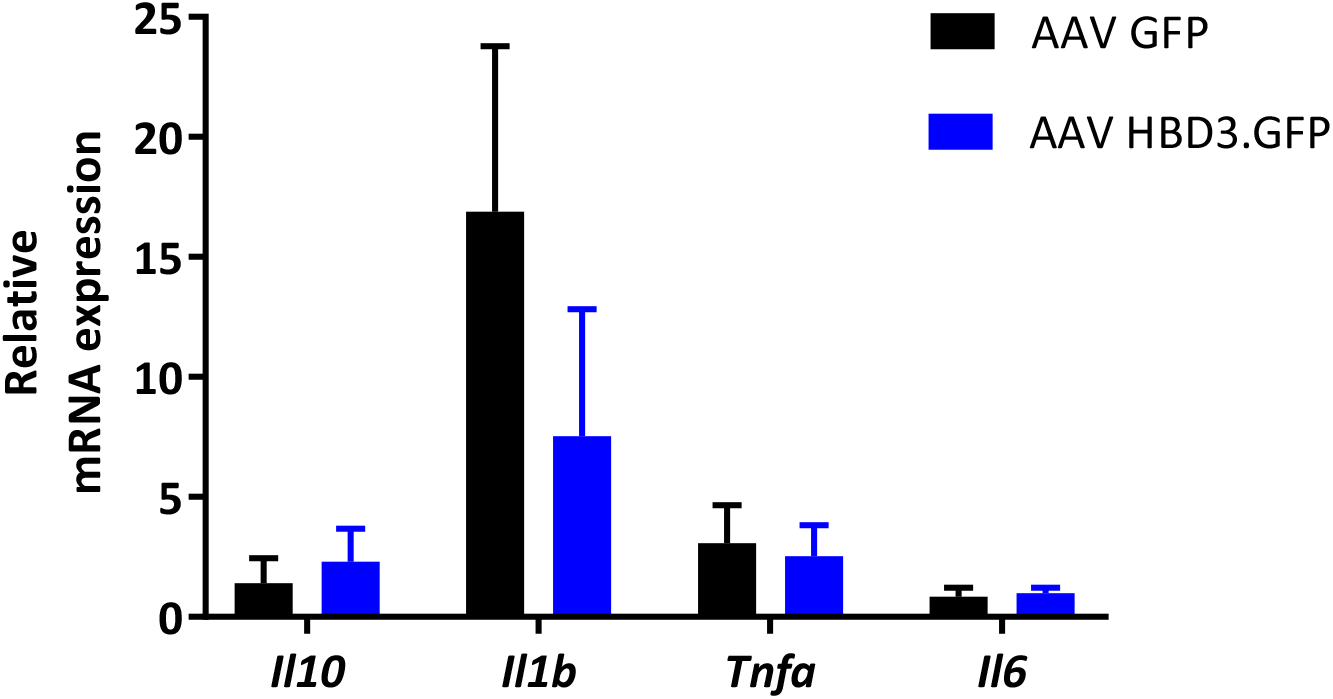
AAV HBD3.GFP does not have an inflammatory cytokine effect on cervical tissue, compared with AAV GFP. Relative mRNA expression was determined in AAV HBD3 and AAV GFP cervices. n=6; data shown as 2^−ΔCT^ data and analysed by two-way ANOVA with post hoc Bonferroni tests.

**Supplementary Table 1:**
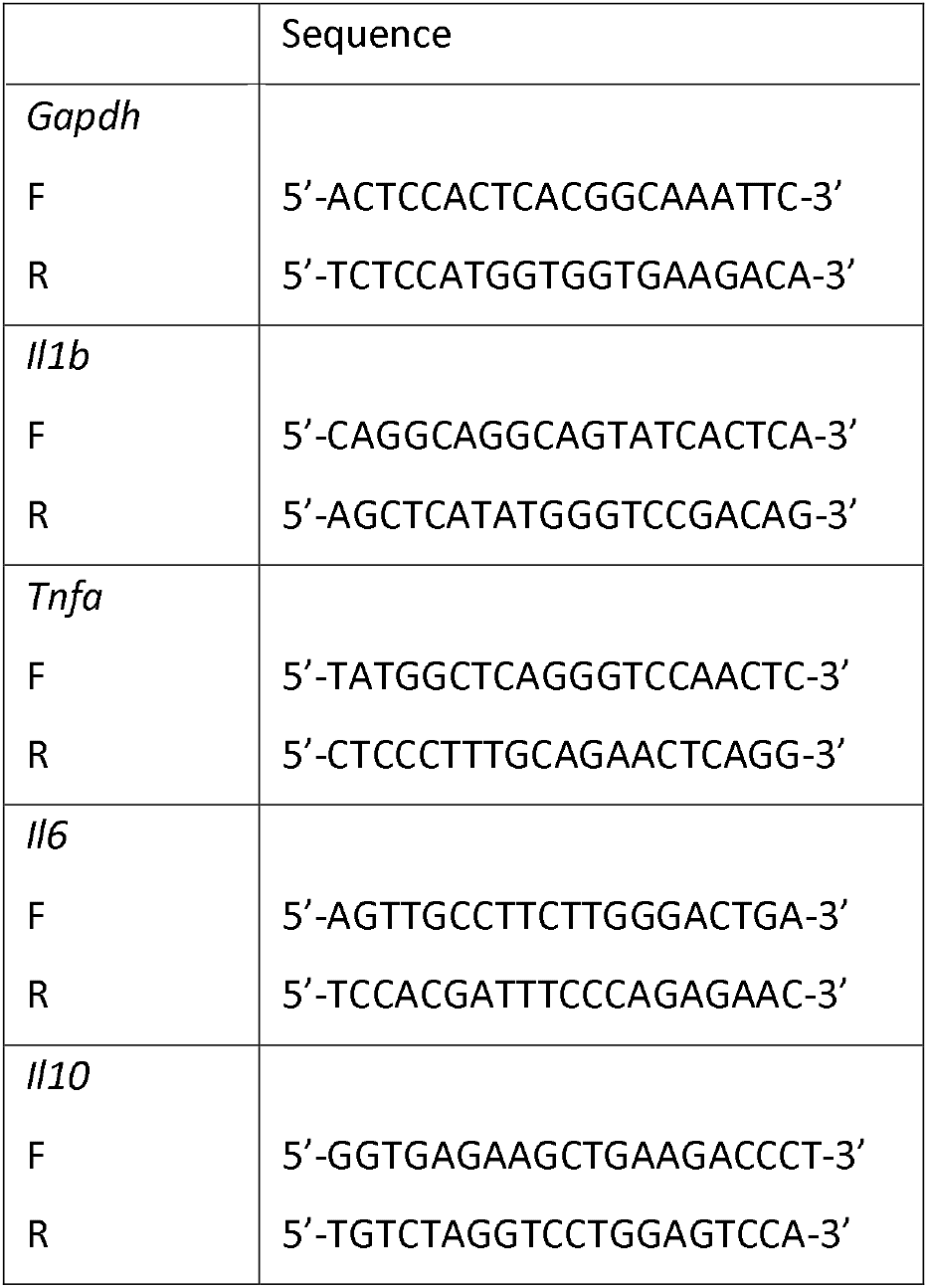
qPCR primer sets

